# Increased Cerebral Blood Flow after single dose of antipsychotics in healthy volunteers depends on dopamine D2 receptor density profiles

**DOI:** 10.1101/336933

**Authors:** Pierluigi Selvaggi, Peter C.T. Hawkins, Ottavia Dipasquale, Gaia Rizzo, Alessandro Bertolino, Juergen Dukart, Fabio Sambataro, Giulio Pergola, Steven C.R. Williams, Federico Turkheimer, Fernando Zelaya, Mattia Veronese, Mitul A. Mehta

**Author notes:** Designated equal contribution as senior author (Mattia Veronese, Mitul A. Mehta). Corresponding author: Pierluigi Selvaggi, Centre for Neuroimaging Sciences, Institute of Psychiatry, Psychology & Neuroscience, King’s College London, De Crespigny Park; SE5 8AF; London, UK.

## Abstract

As a result of neuro-vascular coupling, the functional effects of antipsychotics in human brain have been investigated in both healthy and clinical populations using haemodynamic markers such as regional Cerebral Blood Flow (rCBF). However, the relationship between observed haemodynamic effects and the pharmacological action of these drugs has not been fully established. Here, we analysed MRI-based rCBF data from a placebo-controlled study in healthy volunteers, who received a single dose of three different D2 receptor antagonists and tested the association of the main effects of the drugs on rCBF against normative population maps of D_2_R protein density and gene-expression data. In particular, we correlated CBF changes after antipsychotic administration with non-displaceable binding potential (BP_ND_) template maps of the high affinity D_2_-antagonist Positron Emission Tomography (PET) ligand [^18^F]Fallypride and brain post-mortem microarray mRNA expression data for the *DRD2* gene. For all antipsychotics, rCBF changes were directly proportional to brain D_2_R densities and *DRD2* mRNA expression measures, although PET BP_ND_ spatial profiles explained more variance as compared with mRNA profiles (PET R^2^ range= 0.20-0.60, mRNA PET R^2^ range 0.04-0.20, pairwise-comparisons all p<0.05). In addition, the spatial coupling between ΔCBF and D_2_R profiles varied between the different antipsychotics tested, possibly reflecting differential affinities. Overall, these results indicate that the functional effects of antipsychotics as measured with rCBF are tightly correlated with the distribution of their target receptors in striatal and extra-striatal regions. Our results further demonstrate the link between neurotransmitter targets and haemodynamic changes reinforcing rCBF as a robust in-vivo marker of drug effects. This work is important in bridging the gap between pharmacokinetic and pharmacodynamics of novel and existing compounds.

## Introduction

Antipsychotics are still the preferred choice for the treatment of conditions such as schizophrenia and other mental health disorders with psychotic features (Stroup *et al*, 2009). The main target of most of these compounds is the dopamine D_2_ receptor (D_2_R) (Burt *et al*, 1977; Farde *et al*, 1986) where they act as antagonists or partial agonists. D_2_R occupancy of antipsychotics has been assessed *in vivo* with emission tomography in healthy controls and clinical populations (Agid *et al*, 2007; Kapur *et al*, 1995; Nyberg *et al*, 1995) and it has also been linked with treatment response (Kapur *et al*, 2000). While antipsychotics have been well characterized in terms of their pharmacokinetics (PK) and clinical response, their impact on brain physiology and function is still not well understood. A deeper understanding of these effects is crucial to uncover biological mechanisms driving their clinical efficacy as well as their side effects.

Functional effects of antipsychotics in the brain have been investigated using different neuroimaging tools. Seminal work using Positron Emission Tomography (PET) [^18^F]fluorodeoxyglucose and [^15^O]H_2_O showed that drug naïve first episode psychosis patients, had increased glucose utilization after treatment with antipsychotics (DeLisi *et al*, 1985; Holcomb *et al*, 1996) and greater perfusion (Goozee *et al*, 2014; Miller *et al*, 1997; 2001) in the basal ganglia. Similar results were also obtained in healthy volunteers using a single dose of antipsychotics (Kim *et al*, 2013; Mehta, 2003). Studies using Arterial Spin Labelling (ASL), a Magnetic Resonance Imaging (MRI) sequence designed to quantitatively measure regional cerebral blood flow (rCBF), found results in line with the earlier PET studies. In particular, (Fernández-Seara *et al*, 2011) reported increased rCBF in the striatum and thalamus in healthy volunteers after a single oral dose of 10 mg of metoclopramide (a D_2_R antagonist) and (Handley *et al*, 2013) showed that both haloperidol 3 mg (a D_2_R antagonist) and aripiprazole 10 mg (a D_2_R partial agonist with 5-HT2a antagonism properties) increased rCBF in the striatum in healthy volunteers with a larger effect size for haloperidol. Recently, we tested the effects of a single clinical effective dose of different antipsychotics (Hawkins *et al*, 2018). Consistent with the existing literature, 3 mg haloperidol, 2 mg risperidone and 0.5 mg risperidone increased striatal rCBF as compared with placebo.

One of the major limitations of pharmacological MRI studies stands on the haemodynamic nature of the main functional measures (e.g. BOLD and rCBF). According to the neurovascular coupling model (Attwell and Iadecola, 2002; Logothetis *et al*, 2001) changes in haemodynamic MRI measures reflect a complex cascade of cellular, metabolic and vascular events associated with changes in neuronal activity (Heeger and Ress, 2002; Hoge *et al*, 1999; Singh, 2012). In line with this model, the main effects of drug may be interpreted as the result of dose-dependent enhanced or reduced pre- or post-synaptic activity due to the action of the drug on its targets (Khalili-Mahani *et al*, 2017). Although many antipsychotics bind to numerous receptors, the effects on rCBF have been tacitly attributed to D_2_R blockade. In particular, D_2_R antagonism would lead to enhanced neurotransmitter turnover in the dopaminergic synapses inducing metabolic activity and therefore perfusion demands (Goozee *et al*, 2014; Handley *et al*, 2013). The findings of altered dopamine synthesis capacity after acute antipsychotics administration in human volunteers and rats support this hypothesis (Hertel *et al*, 1996; Ito *et al*, 2009; Vernaleken *et al*, 2008). However, since MRI does not measure neuronal activity directly, making a link between neuro-receptor binding and haemodynamic effects of these compounds requires a degree of conjecture. In addition, in the case of dopaminergic drugs, MRI hemodynamic changes have also been related to non-neuronal mechanisms including the action of D_1_-like receptors on vessels and D_2_-like receptors on perivascular astrocytes (Choi *et al*, 2006; Krimer *et al*, 1998). For these reasons, despite the body of evidence, the neurochemical mechanisms underlying CBF changes after antipsychotics administration remain unclear. The use of multimodal approaches, e.g. combining MR measures with *ex-vivo* autoradiography (Dukart *et al*, 2018), have started to fill this gap of knowledge.

Receptor occupancy theory posits that the magnitude of the drug response is a function of receptor availability (Clark, 1970; Ploeger *et al*, 2009). In other words, incremental changes in functional response correspond to increments of the fraction of receptors bound. This relationship also depends on specific characteristics of each compound, such as receptor affinity (i.e. K_i_). This basic pharmacology principle has been used to characterize the spatial profile of drug effects in the brain (also called “drug fingerprinting”) assuming that brain regions with high density of the target receptor will show higher magnitude of drug effects (Khalili-Mahani *et al*, 2017). This approach has been proposed to describe MRI changes to dopaminergic drugs in preclinical data (Mandeville *et al*, 2013). Indeed, a monotonic increase in rCBF has been observed following injection of an increasing dose of the D_2_/D_3_ antagonist radiotracer [^11^C]raclopride in the striatum of two male rhesus macaques (Sander *et al*, 2013). Interestingly, [^11^C]raclopride non-displaceable binding potential changes (BP_ND_) reflecting changes in D_2_/D_3_ receptor occupancy, correlated with the amplitude of hemodynamic changes (i.e. rCBF): the higher the dose the larger rCBF increase; and in time: BP_ND_ variation and changes in rCBF showed similar temporal profiles. The same group also found an inverse relationship between receptor occupancy and rCBF with the selective D_2_/D_3_ agonist quinpirole in male rhesus macaques (i.e. rCBF decrease in face of dose-dependent increase of receptor occupancy) (Sander *et al*, 2016a). Both studies provide evidence for a neurovascular coupling mechanism linking MR haemodynamic changes and D_2_/D_3_ receptors pharmacological modulation in non-human primates although they are limited to the striatal region.

The aim of the present work is to test in humans whether rCBF changes induced by a single dose of different antipsychotics co-varies with *in vivo* measures of D_2_R distribution. In particular, we employed two datasets from healthy volunteers (Hawkins *et al*, 2018) to evaluate the spatial correlation between rCBF variation in the placebo vs antipsychotic comparison against the population-based receptor density profiles derived from human PET scans using the high affinity D_2_/D_3_ antagonist [^18^F]Fallypride (Mukherjee *et al*, 1995). We also investigated the same relationship at the gene expression level using *post-mortem* mRNA expression measures of *DRD2* (the gene coding for D_2_R) extracted from the Human Allen Brain Atlas (ABA) (Hawrylycz *et al*, 2012a). Brain mRNA expression variation across brain regions has been shown to be associated with resting state fMRI networks suggesting that brain hemodynamic response may be linked to the architecture of the human brain transcriptome (Hawrylycz *et al*, 2015; Richiardi *et al*, 2015). Here, brain microarray mRNA expression data was chosen as a proxy to protein-level receptor density in the human brain. In fact, while post-transcriptional events may alter the relationship between gene expression and protein synthesis (Liu *et al*, 2016), brain mRNA expression maps have been shown to predict *in vivo* proteins level as measured with PET (Beliveau *et al*, 2017; Rizzo *et al*, 2014).

Following the receptor occupancy theory and the neurovascular coupling model proposed by (Sander *et al*, 2013) we hypothesized that there will be a detectable linear relationship between main effects of antipsychotics on CBF measures and D_2_R receptor density profiles evaluated at the protein and gene expression level. Even though brain microarray mRNA expression data from the ABA is noisier and more discrete (i.e. limited number of samples) than the PET BP_ND_ maps, we expect CBF increases after antipsychotics to be linearly associated also with *DRD2* mRNA expression spatial profiles. However, given the fact that mRNA expression only approximates cellular protein levels due to post-transcriptional regulatory mechanisms we predict microarray data to explain less variance in CBF changes than PET derived maps.

## Materials and Methods

### Participants and study design

Data were collected as part of a project approved by the National Research Ethics Service Committee London – Brent (REC reference: 13/LO/1183). Details about participants, protocol and study design have been described in detail in (Hawkins *et al*, 2018). Briefly, forty-two healthy male subjects were enrolled in a double-blind, placebo-controlled, randomised, fully counterbalanced, three-session crossover design. Participants were randomised into two equal parallel study groups (Group 1 age mean/SD 27.6/6.9; Group 2 age mean/SD 28.3/6.3). In the first group participants received placebo, 7.5 mg of olanzapine (OLA), or 3 mg of haloperidol (HAL) on each study day. In the second group, participants received placebo, 0.5 or 2 mg risperidone (lowRIS and highRIS respectively). All but lowRIS dosage were chosen in order to achieve on average at least 60% of D2 receptor occupancy (Kapur *et al*, 2000; Tauscher *et al*, 2004). On dosing day, participants followed a standardised regime. The MRI scan was performed at the time of predicted peak level of plasma concentration of the drug after oral administration (T_max_): approximately 5 hours after drug administration for Group 1 and 2 hours for Group 2 (de Greef *et al.*, 2011; Midha *et al.*, 1989; Nyberg *et al.*, 1997; Tauscher *et al.*, 2002). Study days were seven days apart to allow washout between sessions. After the final visit, a follow-up phone call was made to monitor potential adverse events related to the study drugs.

### MRI acquisition and pre-processing

All scans were conducted on a GE MR750 3 Tesla scanner using a 12-channel head coil. ASL image data were acquired using a 3D pseudo-continuous ASL sequence (3DpCASL) with a multi-shot, segmented 3D stack of axial spirals (8-arms) readout with a resultant spatial resolution (after re-gridding and Fourier Transformation) of 2×2×3mm. Four control-label pairs were used to derive a perfusion weighted difference image. The labelling RF pulse had a duration of 1.5s and a post-labelling delay of 1.5s. The sequence included background suppression for optimum reduction of the static tissue signal. A proton density image was acquired in 48s using the same acquisition parameters in order to compute the CBF map in standard physiological units (ml blood/100g tissue/min). Pre-processing of all CBF data was performed exactly as described in Hawkins *et al*, (2018) (for further details please see Supplementary Materials).

### Receptor density profiles

Figure 1 shows the general framework of the analysis. D_2_R profiles where extracted from an independent [^18^F]Fallypride PET template obtained by averaging six binding potential (BP_ND_) whole brain maps acquired in healthy young volunteers (age range: 18-30 years) who did not participate in the drug study (Dunn *et al*, 2009). [^18^F]Fallypride is a D_2_/D_3_ receptor antagonist and it is a well-established PET radiotracer in the study of D_2_-like receptor distribution in the brain (Mukherjee *et al*, 1995). Compared with other D_2_-like antagonist radiotracers (e.g. [^11^C]raclopride), [^18^F]Fallypride has higher affinity and higher signal-to-noise ratios *in vivo* and therefore provides reliable quantitative measures of D_2_R concentration including extra-striatal brain regions (Mukherjee *et al*, 2002; Stark *et al*, 2018). The [^18^F]Fallypride template was segmented with the Desikan-Killiany Atlas (Desikan *et al*, 2006) and for each of the 85 Regions of Interest (ROIs) of the template, the voxel-wise mean BP_ND_ value was extracted with the *flslmeants* function implemented in the Functional Software Library suite (FSL, FMRIB, Oxford, UK). Conventional parametric modelling of regional BP_ND_ was performed by using the cerebellum as reference region (Ichise *et al*, 2003). Therefore, both left and right cerebellar ROIs (namely “rh_cerebellum_cortex” and “lh_cerebellum_cortex”) were excluded from correlation analyses with CBF profiles. To compare the contribution of striatal vs extra-striatal regions to this association, we also extracted BP_ND_ profiles from a [^11^C]raclopride template obtained from a matchted group of healthy controls (Grecchi *et al*, 2014) (see Supplementary Material).

**Figure 1.**
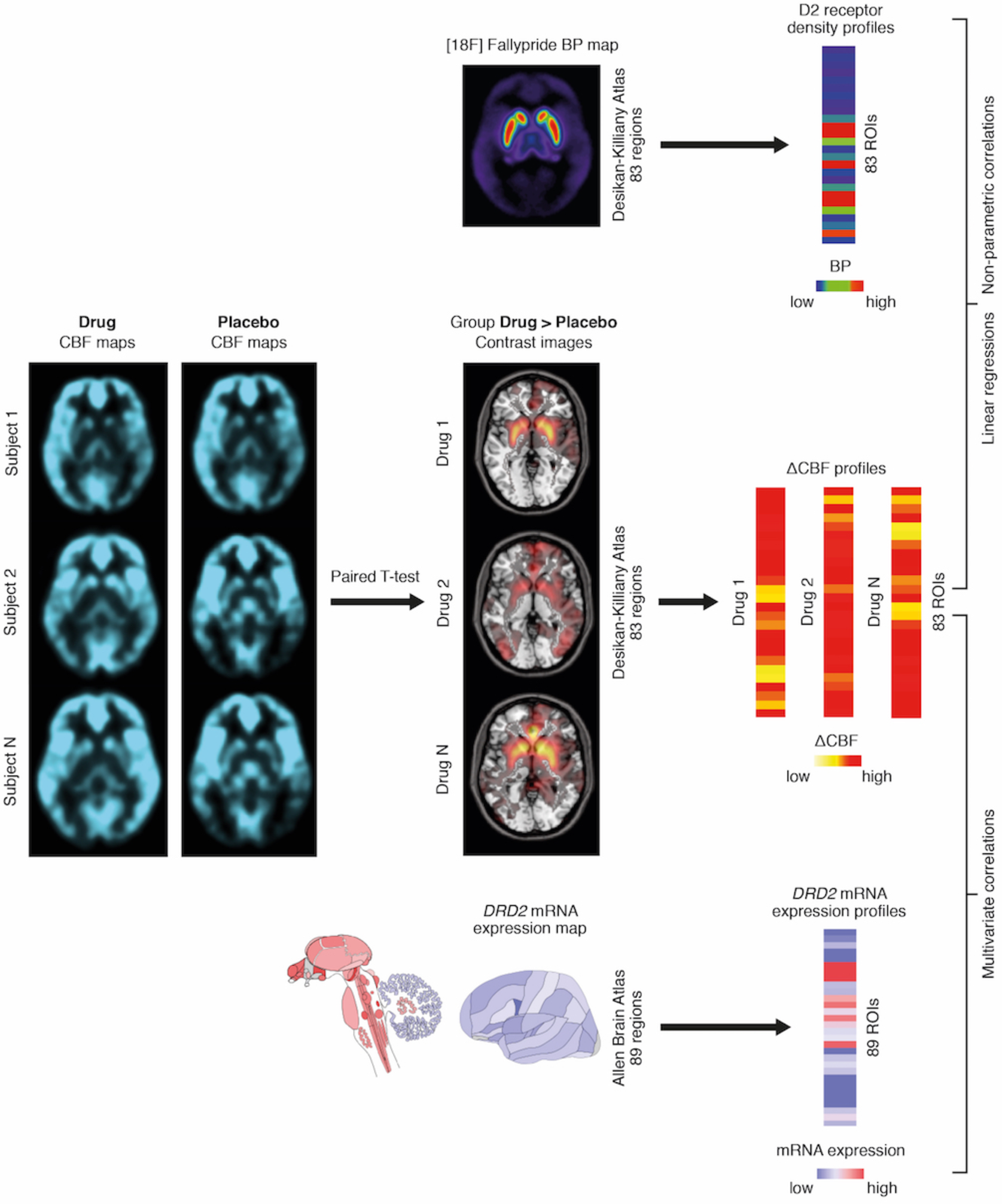
General framework of the analysis. Group-level map showing main effect of drug was computed for each antipsychotic. The resulting maps and the 18F]Fallypride PET template were segmented into 83 ROIs by using the Desikan-Killiany Atlas (Desikan et al, 2006). For each ROI ΔCBF profiles and [18F]Fallypride BPND values were extracted and then correlated. DRD2 gene brain microarray mRNA expression values were extracted from ABA data (http://human.brain-map.org) (Hawrylycz et al, 2012b). MENGA software (http://www.nitrc.org/projects/menga/) (Rizzo et al, 2016) was used to extract DRD2 mRNA expression profiles which then entered multivariate correlation analysis against ΔCBF profiles (please refer to methods section for a more detailed description).

### CBF profiles

CBF profiles were obtained from maps of CBF changes at the group level. In particular, for each antipsychotic, a group-wide paired T-test was performed in SPM12 for each drug vs placebo. Whole brain total blood flow was added as a covariate of no interest in the model to account for peripheral (global) drug effects and between-subjects variability in global brain perfusion (Handley *et al*, 2013; Viviani *et al*, 2009; 2013). In agreement with previous studies (Handley *et al*, 2013; Hawkins *et al*, 2018), we identified statistically significant increases in CBF after drug administration (against placebo) and have thus focussed on this contrast in the remainder of this work. Therefore, DRUG>PLACEBO contrast maps for each antipsychotic where segmented by using the Desikan-Killiany Atlas (Desikan *et al*, 2006) using the same approach used for the extraction of receptor profiles. This resulted in 85 ΔCBF profiles measuring the antipsychotic induced increase of CBF in each ROI. For consistency with the receptor profile data, we carried out correlation analyses excluding the two cerebellum ROIs.

### Statistical analysis for the ΔCBF/receptor density profiles correlations

To test the associations between ΔCBF profiles (derived for each antipsychotic) and D_2_/D_3_ receptor density profiles, linear regression models as implemented in SPSS were used (IBM, SPSS Statistics, Version 23). Normal distribution of the residuals of the regression models were tested by Shapiro-Wilk test. BP_ND_ data were transformed using a natural logarithmic function (ln) so that the residuals conformed to a normal distribution. For all linear regression models Mahalanobis distance and Cook’s distance were computed in order to explore the presence of multivariate outliers and estimate the presence of highly influential data points. To identify multivariate outliers, Mahalanobis distance values were compared to a chi-square distribution with degrees of freedom equal to the number of variables (two in this case) with p= 0.001 (Finch, 2012; Tabachnick and Fidell, 2013). Any data point with Cook’s distance higher than 1 was considered as highly influential outlier and excluded from the analysis (Cook and Weisberg, 1982). Non-parametric Spearman’s correlations between ΔCBF profiles and receptor BP_ND_ profiles were also performed as a countercheck. In addition, Fisher’s r-to-z transformation was performed to test pairwise significance of the difference between correlation coefficients of ΔCBF profiles between different antipsychotics. Asymptotic covariance method was adopted to account for the fact that correlations had one variable in common (Lee and Preacher, 2013)

### mRNA profiles and genetic correlations

*DRD2* gene brain microarray mRNA expression values were extracted from ABA data (http://human.brain-map.org) by using the Multimodal Environment for Neuroimaging and Genomic Analysis (MENGA) toolbox (http://www.nitrc.org/projects/menga/) (Rizzo *et al*, 2016). The same toolbox was used to carry out correlations with ΔCBF profiles of each antipsychotic. First, antipsychotics’ contrast images (DRUG>PLACEBO) were resampled in the ABA space. Then, each CBF image sample was spatially matched with the corresponding genomic ABA sample within a search window of a sphere of 5mm radius centred on the MNI coordinates of the ABA sample. Both CBF contrast image and ABA data were then segmented using the list of structures (N= 169) provided by the ABA (Hawrylycz *et al*, 2012b). A subset of 89 structures (ROIs), each containing at least one genomic sample for all the six ABA brains (donors), was selected to perform correlations between ΔCBF profiles and gene expression. After completing the matching and the extraction of ΔCBF and gene expression profiles, two different correlations were performed: 1) between-donors correlation or gene auto-correlation returning the biological variability of the spatial profile of mRNA expression between donors (the higher the gene autocorrelation the lower the heterogeneity in mRNA expression spatial profile between donors); 2) correlation between each gene expression and the ΔCBF by ROIs also called cross-correlation. A Principal Component Analysis (PCA) on mRNA expression measures of the 6 ABA donors was performed beforehand in order to extract the component that accounted at least for the 95% of the total variance in the mRNA expression data. In particular, the PCA was performed on an 89−6 matrix (89 ROIs by 6 donors) and represented a consistent spatial mRNA expression profile across all donors. This component was then used in the regression model against CBF profiles. Significance was assessed with a bootstrapping approach resulting in a chance likelihood of the correlation coefficient expressed as a %. More specifically, ROIs were permuted within donors repeating the correlation between PCA component and CBF profiles 1,000 times, in order to obtain a measure of the likelihood that the correlation found was different from chance level. A multivariate spatial correlation using MENGA was also performed between *DRD2* gene expression profiles and [^18^F]Fallypride BP_ND_ maps. As for protein density profile analysis, Fisher’s r-to-z transformation was performed to test pairwise significance of the difference between correlation coefficients of ΔCBF profiles between different antipsychotics and also between PET and mRNA profiles.

## Results

### CBF changes with D_2_ receptor profiles correlations

All antipsychotic ΔCBF profiles significantly correlated with [^18^F]Fallypride BP_ND_ template values (Table 1 and Figure 2). HAL ΔCBF had the strongest correlation with [^18^F]Fallypride BP_ND_ (R_linear_= 0.78) followed by RIS (R_linear_= 0.73 and R_linear_= 0.72 for lowRIS and highRIS respectively) and OLA (R_linear_= 0.48). Results from linear models and non-parametric Spearman correlations were consistent. In all linear regressions, none of the data points were identified as a highly influential outlier (all Cook’s distances> 1) or multivariate outliers (all Mahalanobis distances p>0.01). The rank of order and R_linear_ values matched the variation in affinity with D_2_ receptor (McCormick *et al*, 2010), with the stronger the association between ΔCBF profiles and D_2_ receptor densities the lower the Ki (Table 1 and Figure S2). In the pairwise correlation comparisons we found significant difference for HAL vs OLA (z= 5.46), lowRIS vs OLA (z= 4.19) and vs highRIS vs OLA (z= 3.90) (all p_two-tailed_ < 0.01, Bonferroni corrected, Figure S2). All the other comparisons were not significant. To compare striatal vs extra-striatal contributions to this association we also performed correlations between ΔCBF and receptor density profiles by extracting BP_ND_ values from a [^11^C]raclopride PET template. We found weaker correlation with [^11^C]raclopride BP_ND_ as compared with [^18^F]Fallypride BP_ND_ (see Supplementary Material).

**Table 1.** Summary of the correlations of ΔCBF profiles D_2_ receptor profiles ([^18^F]Fallypride BP_ND_) and of genomic multivariate cross-correlations of antipsychotics’ ΔCBF profiles with *DRD2* mRNA expression profiles.

| **DRUG>PLA**<br>**contrast** | **[<sup>18</sup>F]Fallypride BP<sub>ND</sub>** |  |  |  | ***DRD2* mRNA**<br>**expression** |  |
| --- | --- | --- | --- | --- | --- | --- |
|  | **R<sub>linear</sub>** | **p<sub>linear</sub>** | **Spearman rho** | **p<sub>Spearman</sub>** | **R** | **Chance**<br>**likelihood** |
| HAL | + 0.78 | p<0.001 | 0.61 | p<0.001 | + 0.34 | 0 % |
| OLA | + 0.48 | p<0.001 | 0.51 | p<0.001 | + 0.21 | 0 % |
| lowRIS | + 0.73 | p<0.001 | 0.76 | p<0.001 | + 0.43 | 0 % |
| highRIS | + 0.72 | p<0.001 | 0.61 | p<0.001 | + 0.45 | 2 % |

**Figure 2.**
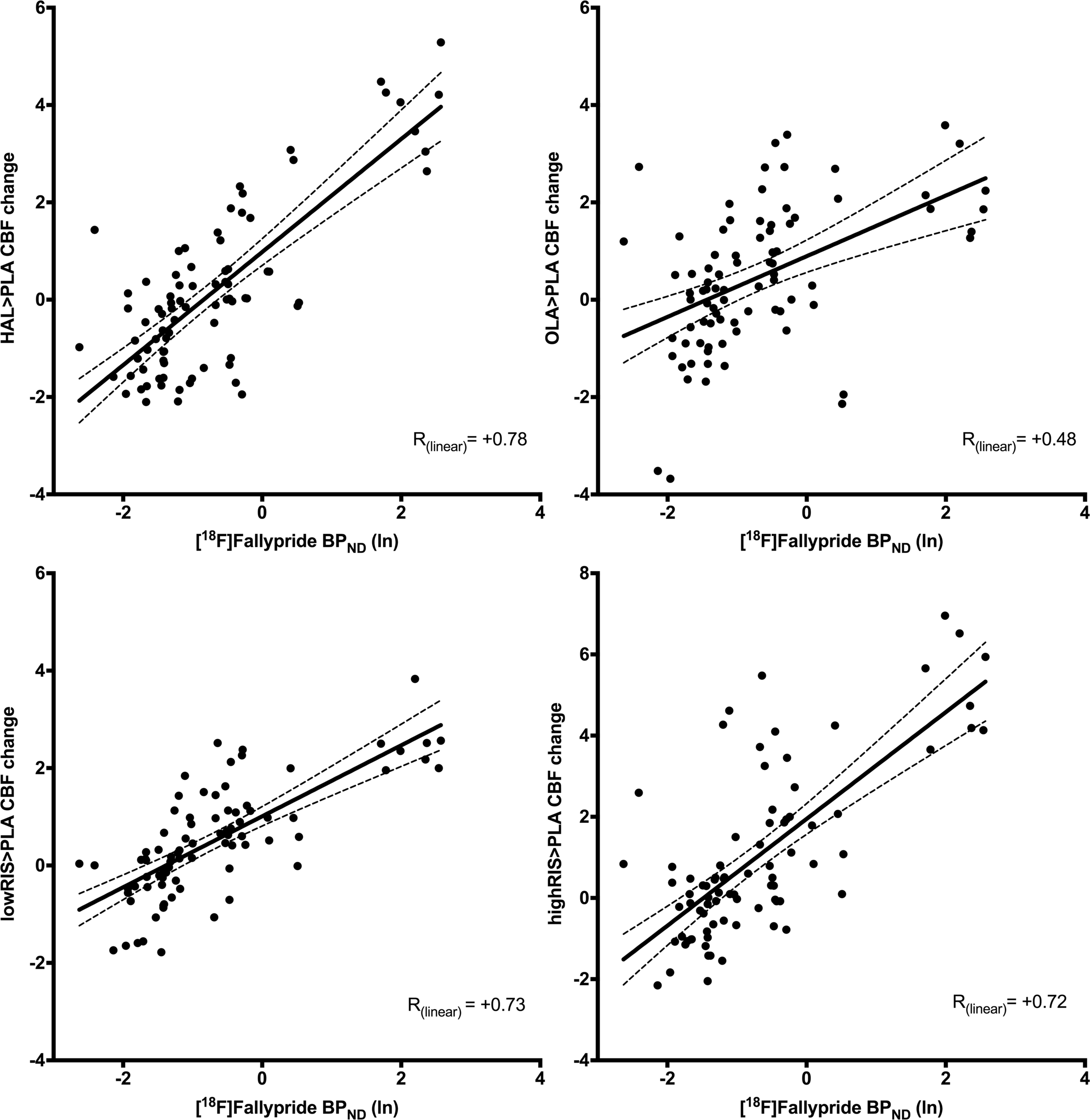
Scatterplots of ΔCBF/receptor density profiles correlations. Top row: scatterplot of the correlation between HAL ΔCBF profiles and [18F]Fallypride BPND (left) and of the correlation between OLA ΔCBF profiles and [18F]Fallypride BPND (right). Bottom row: scatterplot of the correlation between lowRIS ΔCBF profiles and [18F]Fallypride BPND (left) and of the correlation between highRIS ΔCBF profiles and [18F]Fallypride BPND (right). Dashed lines indicate 95% confidence bands.

### mRNA expression correlations

The average correlation coefficient (R^2^) of the genomic autocorrelation analysis for the *DRD2* gene was 0.575 (standard deviation= 0.058) for the six donors. This result indicated good stability between donors of *DRD2* mRNA expression spatial profile (Rizzo *et al*, 2016). For the different antipsychotic drugs, the correlation coefficients were all positive and statistically significant (Figure 3 and Table 1) and also significantly lower than those obtained for the PET template (PET R^2^ range= 0.20-0.60; mRNA PET R^2^ range 0.04-0.20; pairwise-comparisons all p<0.05). As for the correlation with [^18^F]Fallypride values, genomic mRNA expression correlations qualitatively mirrored Ki differences (McCormick *et al*, 2010) between antipsychotics at D_2_R (Figure S2). However, none of the pairwise comparisons between correlation coefficients between antipsychotics were statistically significant after correction for multiple comparisons.

**Figure 3.**
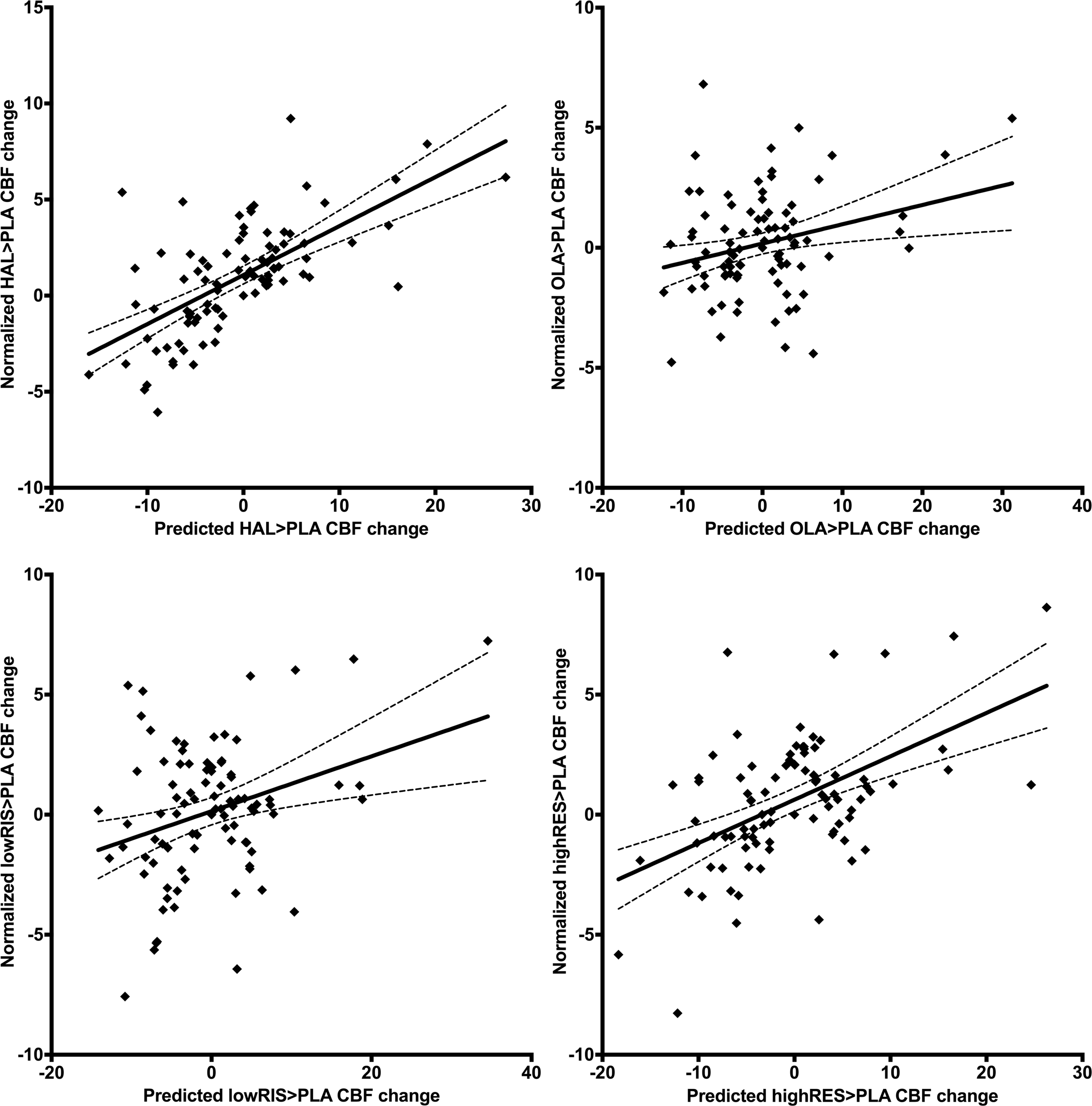
Scatterplots of the genomic correlation. For all scatterplots on the y axis normalized DRUG>PLACEBO CBF changes and on the x axis DRUG>PLACEBO CBF changes predicted by the first Principal Component of the mRNA expression measures of the 6 ABA donors. Top right: HAL; Top left: OLA; Bottom right: lowRIS; Bottom left: highRIS. Dashed lines indicate 95% confidence bands.

## Discussion

The aim of the present study was to investigate the relationship between the effects of single clinical effective doses of antipsychotics on rCBF and receptor distribution profiles in the brain as indexed by [^18^F]Fallypride BP_ND_ values and brain *DRD2* mRNA expression profiles. Consistently with our hypothesis, we found that for all compounds there was a spatial coupling between drug-induced CBF changes and D_2_ receptor density profiles (at both protein and gene expression level). In addition, we found that mRNA data explained less variance in CBF changes than PET derived map.

### Receptor density profiles

The association between CBF changes induced by all antipsychotics and receptor brain spatial distribution of D_2_R matches earlier evidence in non-human primates showing large CBF increases after injection of the D_2_ antagonist PET tracer [^11^C]raclopride in brain regions with high D_2_R density (Sander *et al*, 2013). This suggests that the relationship between the physiological response to D_2_R antagonist and D_2_R availability described by (Sander *et al*, 2013) in preclinical models also exists *in vivo* in humans. Furthermore, we have shown that the relationship between ASL-CBF increases after antipsychotic administration also matched [^18^F]Fallypride BP_ND_ values in extra-striatal ROIs significantly populated by D_2_R such as the thalamus and the amygdala, even though they show lower BP_ND_ values as compared with striatal ROIs. These results extend earlier evidence (Sander *et al*, 2013; 2016b) and suggest that the linear coupling between CBF response to dopaminergic drugs and D_2_R concentration might also be a valid model outside the striatum. This interpretation is also supported by the weaker correlation of ΔCBF with [^11^C]raclopride BP_ND_ as compared with [^18^F]Fallypride BP_ND_ (Supplementary Material).

### Microarray mRNA expression data

We found that for all antipsychotics, the ΔCBF profiles also correlated with microarray mRNA expression data extracted from the ABA. All genomic correlations were positive and therefore in the same direction of [^18^F]Fallypride BP_ND_ linear models. Notably, microarray mRNA expression measures explained less variance in ΔCBF than [^18^F]Fallypride BP_ND_ values. This is consistent with our hypothesis motivated by the existence of a large variability between mRNA expression and protein synthesis due to post-transcriptional regulation mechanisms (Liu *et al*, 2016). Our findings are consistent with previous works that have linked the spatial architecture of the brain transcriptome to brain structure (Grecchi *et al*, 2017; Veronese *et al*, 2015), function (Hawrylycz *et al*, 2015; Richiardi *et al*, 2015) and *in vivo* measures of brain proteins (Gryglewski *et al*, 2018; Rizzo *et al*, 2014; 2016; Veronese *et al*, 2016a). Here we could show that *post mortem* brain mRNA expression data may potentially be used to map also variations in MRI-based functional response to drug stimulation. However, such mapping has still many limitations to be addressed in order to adopt it extensively in the context of pharmacological-MRI studies. For instance, it might be difficult to use mRNA expression mapping in neurotransmitter systems where protein synthesis is highly dependent on post-transcriptional regulation mechanisms (e.g. serotonin system) (Beliveau *et al*, 2017; Rizzo *et al*, 2014). While further studies are needed to fully validate this mRNA expression-MRI approach, it might be especially valuable for profiling the functional effects of drugs with poorly characterized or unknown targets. Interestingly, as for the correlation with receptor density profiles, the rank order of correlations between ΔCBF and *DRD2* mRNA expression measures mirrored affinity profiles of the compounds, although these differences were only numerical. It is worth noting that the variance explained by genomic correlation was lower than the PET correlations (Table 1), which might have reduced the chance of detecting any significant difference on the pairwise comparison between the correlation coefficients.

These findings represent a further evidence supporting the hypothesized PK/PD model of antipsychotic effect of CBF measures (Mandeville *et al*, 2013; Sander *et al*, 2013). Indeed, we were able to show that quantitative measures of brain functional effect of antipsychotics (i.e. CBF changes) are directly associated with receptor density measures evaluated at different scales: mRNA expression and protein density.

### Differential strength of association between ΔCBF and receptor density profiles

The correlation strength between ΔCBF and receptor density measured with PET varied between the different antipsychotics tested. One possible interpretation of this difference might be related to the differential affinities for these compounds to D_2_R (Dukart *et al*, 2018). Our data seems to be in line with this hypothesis. In fact, the strengths of the association were higher for antipsychotics with higher affinity for D_2_R (Table 1 and Figure S1). In particular, HAL was the drug with the highest correlation coefficient and the lowest Ki, whereas OLA showed the lowest correlation coefficient and the highest Ki. Both lowRIS and highRIS were in the middle between HAL and OLA. Another possible interpretation might be that related with the different secondary affinities between the drugs. For instance, for compounds with an high affinity with 5HT2a receptors like risperidone and olanzapine, part of the effect on CBF might also be linked with a mechanism different from D_2_R blockade (Goozee *et al*, 2014). Nonetheless, differences in brain disposition of antipsychotics have been reported in previous studies (Kornhuber *et al*, 2006; Rodda *et al*, 2006). In particular, animal data suggested that antipsychotics (including the ones considered in the present work) show different blood-brain barrier penetration and brain clearance leading to dissimilar spatial distribution in the brain (Loryan *et al*, 2016). Even though these dissimilarities have been reported to be only moderate (Loryan *et al*, 2016), brain disposition is a factor to be considered in addition to receptor affinity when linking the pharmacodynamics of each antipsychotic with receptor occupancy. Therefore, without maps of brain deposition variation, conclusions regarding the D_2_R affinities and strength of associations between receptors and CBF are necessarily incomplete.

### Limitations

A number of limitations need to be considered for the present study. First, we used population-based profiles of D_2_R density as the spatial architecture of D_2_R is typically consistent across individuals (Rizzo *et al*, 2014; Veronese *et al*, 2016b). For example, the striatum always has higher D_2_R density than the thalamus which in turn has higher D_2_R density than the cortex. However, variance in density between individuals within the same brain region (Farde *et al*, 1995) may drive inter-individual differences in the drug functional response. Individualized receptor profiles mapping is therefore necessary to bring more precision to the method and to further validate the present findings. Nevertheless, normative atlases for protein and mRNA expressions have proven useful in many different applications, suggesting that the core *spatial* architecture of the brain receptor systems is consistent across individuals (Beliveau *et al*, 2017; Rizzo *et al*, 2014; 2016; Veronese *et al*, 2016b).

In addition, we considered only local effects by matching CBF changes and D_2_R receptor density measures with the same ROIs. However, studies in patients and healthy volunteers showed that antipsychotics also produce changes in functional connectivity in rs-fMRI suggesting the existence of downstream effects (Cole *et al*, 2013; Sarpal *et al*, 2015) that were not included in our analyses.

As discussed above, increases in CBF after acute antipsychotic challenge have been usually interpreted as the result of a neuronal metabolic changes due to D_2_R blockade. However, D_2_Rs are also present in perivascular astrocytes modulating brain hemodynamic changes (Choi *et al*, 2006). In addition, both risperidone and olanzapine, but not haloperidol, act as antagonists at the serotonin-2a receptors (5HT2a) (Meltzer, 1999). Blockade of 5HT2a receptors on smooth-muscle cells of brain arteries has been shown to induce relative vasodilation in animal models (i.e. blocking vasoconstriction caused by the endogenous ligand serotonin) (Kovács *et al*, 2012). Therefore, the degree to which these non-neuronal effects contribute to the associations described here is not known.

## Conclusion

Understanding the link between neurochemical changes and brain function is crucial to uncover mechanisms underlining the effects of psychopharmacological treatment and between-subjects variability in response. In this work we investigated the case of antipsychotics whose functional effect, evaluated as changes in CBF measures, mirror the well-known spatial distributions of their main target (i.e. D_2_R). The characterisation of the haemodynamic response using this multimodal approach might be applicable also for other classes of drugs and it becomes particularly valuable for profiling compounds known to bind to multiple targets or with unknown or poorly characterised targets. Finally, our work demonstrates that the use of MRI 3D pCASL as a measure of regional CBF offers an efficient, non-invasive tool to investigate CNS penetration in the process of psychotropic drug development that is also linked with brain chemistry.

### Funding and Disclosure

Contract grant sponsor: Hoffmann – LaRoche Pharmaceuticals.

This paper represents independent research part funded by the National Institute for Health Research (NIHR) Biomedical Research Centre at South London and Maudsley NHS Foundation Trust and King’s College London that support PS, OD, SCRW, FZ, MV and MAM. The views expressed are those of the authors and not necessarily those of the NHS, the NIHR or the Department of Health and Social Care. PS is supported by a PhD studentship jointly funded by the NIHR-BRC at SLaM and the Department of Neuroimaging, King’s College London.

AB is a stockholder of Hoffmann-La Roche Ltd. He has also received consulting fees from Biogen and lecture fees from Otsuka, Janssen, Lundbeck. FS is a former employee of F. Hoffmann-La Roche Ltd.GP has been the academic supervisor of a Roche collaboration grant (years 2015-16) that funds his salary. JD is current employees of F. Hoffmann-La Roche Ltd. and received support in form of salaries. SCRW has received grant funding from the Medical Research Council (UK), Wellcome Trust (UK), National Institute for Health Research (UK) and support for investigator led studies from Takeda, Pfizer, Lundbeck, P1Vital, Roche and Eli Lilly. In the past 3 years MAM has acted as an advisory board member for Lundbeck and Forum Pharmaceuticals. He also holds research funding from Lundbeck, Takeda and Johnson & Johnson. No other conflict of interested are disclosed.

## Acknowledgments

MRI data were collected at the NIHR/Wellcome Trust King’s Clinical Research Facility (CRF). Authors are grateful to the CRF team and to Dr Ndabezinhle Mazibuko and Stephanie Stephenson (Department of Neuroimaging, Institute of Psychiatry, Psychology and Neuroscience, King’s College London, London, United Kingdom) for their help on data acquisition. We also gratefully acknowledge the work of Dr Joel Dunn (Kings College London and Guy’s and St Thomas’ PET Centre, School of Biomedical Engineering and Imaging Sciences, King’s College London, London, UK) who contributed to the creation of PET templates. We also thank Gil Brown (London College of Communication, UAL London, UK) for her help on illustrations. Finally, we thank all the volunteers who took part in the study.

Supplementary information is available at the *Neuropsychopharmacology* website

